# Heritable and climatic sources of variation in juvenile tree growth in an Austrian common garden experiment of Central European Norway spruce populations

**DOI:** 10.1101/2022.01.10.475611

**Authors:** Laura Morales, Kelly Swarts

## Abstract

We leveraged publicly available data on juvenile tree height of 299 Central European Norway spruce populations grown in a common garden experiment across 24 diverse trial locations in Austria and weather data from the trial locations and population provenances to parse the heritable and climatic components of juvenile tree height variation. Principal component analysis of geospatial and weather variables demonstrated high interannual variation among trial environments, largely driven by differences in precipitation, and separation of population provenances based on altitude, temperature, and snowfall. Tree height was highly heritable and modeling the covariance between populations and trial environments based on climatic data led to more stable estimation of heritability and population × environment variance. Climatic similarity among population provenances was highly predictive of population × environment estimates for tree height.

## Introduction

Under rapid anthropogenic climate change, ecosystem dominant conifer species, such as Norway spruce (*Picea abies*), are facing environmental conditions to which they have not been adapted (Lindner et al. 2010; Seidl et al. 2017). Although Norway spruce has broad genetic diversity and geographic adaptation, long generation times and the confounded effects of population structure and local adaptation make breeding populations for new environments challenging (Leslie et al. 2012; Yeaman et al. 2016; Wang et al. 2020). Common garden experiments can be used to evaluate a select number of diverse individuals or populations in different environments (Oleksyn et al. 1998; Kapeller et al. 2012; Aitken and Bemmels 2016). Assisted migration can then be applied, where well-adapted individuals are introduced into locations with compatible (current or forecasted) climates (Williams and Dumroese 2013; Aitken and Bemmels 2016).

Precise estimation of genetic and environmental effects on tree growth phenotypes is critical for assisted migration and breeding program selections (Williams and Dumroese 2013; Aitken and Bemmels 2016). In addition to proper experimental design, germplasm choice, and phenotyping methods, statistical approaches can also increase the genetic signal and prediction accuracy of phenotypes (Heslot et al. 2015; Crossa et al. 2017; Isidro y Sánchez and Akdemir 2021). For example, leveraging genetic marker, geospatial, and/or climatic data to model relationships between individuals, trial locations, and genotype × environment interactions is a common approach in agricultural and forest genetics and breeding (Heslot et al. 2014; Crossa et al. 2016; Rodríguez-Álvarez et al. 2018; Avanzi et al. 2019; Bustos-Korts et al. 2019).

In this study, we combined publicly available datasets on (a) juvenile tree height from a common garden experiment of Norway spruce, in which 299 Central European populations were evaluated across five years in 24 locations in Austria and (b) historical monthly precipitation and temperature measurements from the common garden trial locations and population provenances (Efthymiadis et al. 2006; Hiebl et al. 2009; Chimani et al. 2011, 2013; Kapeller et al. 2012, 2017). Previous studies have used this dataset to understand varying selection and intraspecific variation in climate response from an ecological perspective (Kapeller et al. 2012, 2017). Here, we employed agricultural statistical methods to parse and to predict the heritable (genetic) and environmental components of phenotypic variance in tree height.

## Materials and methods

### Common garden experiment

We used a publicly available dataset to investigate population and environmental effects on juvenile tree growth in Norway spruce (*Picea abies*) (Nather and Holzer 1979; Schulze 1985; Kapeller et al. 2012, 2016, 2017). The plant material was derived from 299 Norway spruce populations collected across Central Europe during commercial seed harvests in 1971. Seeds from each population were germinated and seedlings were then transplanted into a nursery in Austria in 1973. In 1978, a total of 65,534 five-year-old trees were transplanted into 24 trial locations across Austria in a balanced incomplete block design. On average, each population was grown at two locations, 27 populations were grown in each location, and 102 trees (replications) from each population were split into three randomized blocks within each location. The height (cm) of each tree was recorded at the ages of 7-10 and 15 years in 1980-1983 and 1988, respectively, resulting in a total of 300,310 observations. The previously published dataset also included the altitude, latitude, and longitude for all trial locations and for 278 population provenances.

### Weather data

We accessed monthly precipitation (total, liquid, solid) and temperature data from the publicly available Historical Instrumental Climatological Surface Time Series of the Greater Alpine Region (HISTALP) resource (Efthymiadis et al. 2006; Hiebl et al. 2009; Chimani et al. 2011, 2013). The HISTALP dataset includes monthly temperature and precipitation grids from 1780-2009 and 1801-2003, respectively, at 5 min x 5 min resolution ranging from 4-19°E and 43-46°N. We extracted data from the original Network Common Data Form (NetCDF) formatted HISTALP files using the “netcdf4” package in R (R Core Team 2020; Pierce 2021).

### Climatic analysis

All altitude, latitude, longitude, and HISTALP weather variables used in the analyses described in this subsection were centered and scaled using the “scale” function in R (R Core Team 2020). We conducted principal component analysis (PCA) using the “FactoMineR” package in R (Lê et al. 2008; R Core Team 2020). To investigate spatiotemporal variation across the common garden experiment, we extracted monthly temperature and precipitation data from the grids closest to the trial locations in 1980-1983 and 1988 from the HISTALP dataset. We then conducted PCA using the altitude, latitude, longitude and monthly weather data from each trial location in each year, henceforth referred to as trial environments (PCA_Env_). To explore climatic differences among population provenances, we extracted monthly temperature and precipitation data from the grids closest to the population provenances from 1801 (earliest year with both temperature and precipitation data in HISTALP) to 1970 (year before seed harvests). Weather data from this time period should reflect the climates experienced by the parental populations of the trees evaluated in the common garden experiment. We then conducted a PCA using the altitude, latitude, longitude, and monthly weather data from each population provenance (PCA_Prov_).

We used the altitude, latitude, longitude, and HISTALP weather data described above to model the relationships among population provenances and environments (trial location-years). We first calculated Euclidian distance matrices using said geospatial and weather variables for all pairs of population provenances (***D***_*Prov*_) and trial environments (***D***_*Env*_) with the “dist” function in R (R Core Team 2020). We then transformed the distance matrices into similarity matrices as 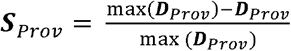 and 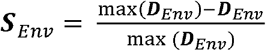 for further use in phenotypic modeling.

### Phenotypic analysis using climatic data

Due to the large number of observations (N=300,310) in the common garden experiment, it was not computationally feasible to specify complex variance-covariance matrices for the entire dataset in one model. As such, we used a two-stage approach common in multi-environment plant breeding and genetics studies (Piepho et al. 2012, 2020), with mixed linear models fit with the “breedR” package in R (Muñoz and Sanchez 2020; R Core Team 2020). In the first stage, we fit the following mixed model within each environment (location-year):

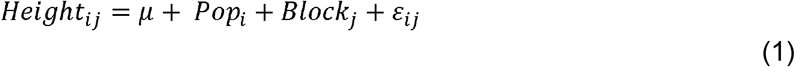

where *Height*_*ij*_ is the tree height response, *µ* is the overall mean, *pop*_*i*_ is the fixed effect of population *i, Block*_*j*_, is the random effect of block *j*, and *ε*_*ij*_ is the random error. The random effects were independent and identically normally distributed as 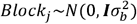 and 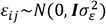. We then extracted the fitted values (*ŷ*_*ij*_) for each population *i* within each block *j* from each within-environment model (1) as

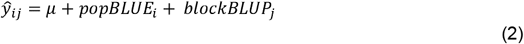

Where *µ* is the intercept, *popBLUE*_*i*_ is the best linear unbiased estimate (BLUE) of population *i*, and *blockBLUP*_*j*_ is the best linear unbiased predictor (BLUP) of block *j*. We combined the fitted values across environments, resulting in a reduced and refined set of phenotypes for further analysis (N = 7,314). Only fitted values for populations with climatic information were included in the second stage of the analysis (278/299 populations).

In the second stage, we fit a naïve across-year mixed model to understand the environmental and genetic (population-level) effects on juvenile tree growth as:

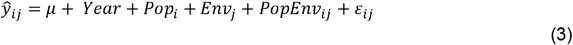

where *ŷ*_*ij*_ are the fitted values from each within-environment model (2), *µ* is the overall mean, *Year* is the continuous fixed effect of year, *Prov*_*i*_ is the random effect of population *i, Env*_*j*_ is the random effect of trial environment *j, ProvEnv*_*ij*_ is the random effect of the interaction between population *i* and trial environment *j*, and *ε*_*ij*_ is the random error. The random effects were assumed to be independent and identically normally distributed, where 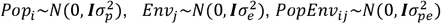, and 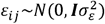.

We also fit an across-year mixed model with form similar to model (3) using historical weather data to model similarity among trial environments and populations provenances as:

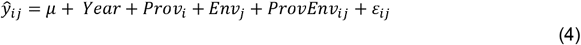

where *ŷ*_*ij*_ are the fitted values from each within-environment model (2), *µ* is the overall mean, *Year* is the continuous fixed effect of year, *Prov*_*i*_ is the random effect of population provenance *i, Env*_*j*_ is the random effect of trial environment *j, provEnv*_*ij*_ is the random effect of the interaction between population provenance *i* and trial environment *j*, and *ε*_*ij*_ is the random error. The random effects were distributed as 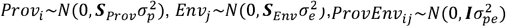, and 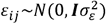.

We fit naïve within-year mixed models with the form in order to understand the environmental and genetic effects on juvenile tree height within developmental stage (tree age) as:

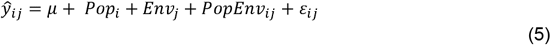

where *ŷ*_*ij*_ are the fitted values from each within-environment model (2) from a given year (1980, 1981, 1982, or 1983), *µ* is the overall mean, *Pop*_*i*_ is the random effect of population *i, Env*_*j*_ is the random effect of trial environment *j, PopEnv*_*ij*_ is the random effect of the interaction between population and trial environment *j*, and *ε*_*ij*_ is the random error. The random effects were assumed to be independent and identically normally distributed, where 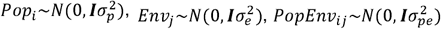, and 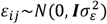.

We also fit a within-year mixed model with form similar to model (5) using environmental and population provenance weather data as:

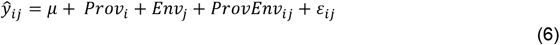

where *ŷ*_*ij*_ are the fitted values from each within-environment model (2) from a given year (1980, 1981, 1982, or 1983), *µ* is the overall mean, *Prov*_*i*_ is the random effect of population provenance *i, Env*_*j*_ is the random effect of trial environment *j, ProvEnv*_*ij*_ is the random effect of the interaction between population provenance *i* and trial environment *j*, and *ε*_*ij*_ is the random error. The random effects were distributed as 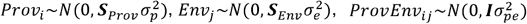, and 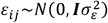, where ***S***_*Prov*_ and ***S***_*Env*_ were subset for the population provenances and environments included in each within-year model, respectively.

In models (4) and (6), we used the term “population provenance” rather than “population” for two reasons: (i) to distinguish models (4) and (6) from models (3) and (5) and (ii) the population variance was modeled as the similarity matrix estimated from climatic data from the provenances (seed source locations) of the populations. Because of insufficient population × environment replication in 1988, we did not fit within-year models for 1988.

For each second stage model (3-6), we estimated broad-sense heritability (*H*^2^) as

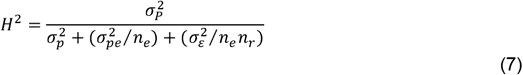

where *n*_*e*_ and *n*_*r*_ are the number of trial environments (*n*_*e*_ = 10 across years; *n*_*e*_ = 2 within years) and replications per population/provenance per trial environment (*n*_*r*_ = 3), respectively, and 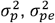, and 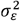 are the variance components for population provenance, population/provenance × environment, and error, respectively (Holland et al. 2003; Piepho and Moehring 2007).

### Predictions using climatic covariance

Genomic prediction, in which phenotypes are predicted based on genetic similarity (kinship), is a widely used approach in plant and animal breeding for quantitative traits (controlled by many genes with small individual effects) (Hickey et al. 2017). In addition, genotype × environment interactions can be modeled with both genetic and weather-based covariance structures to improve genomic prediction accuracy (Jarquín et al. 2014; Chen et al. 2017; Gillberg et al. 2019; Costa-Neto et al. 2021). We extracted the population, environment, and population × environment BLUPs from the naïve across-year phenotypic model (3) and then fit the following mixed models:

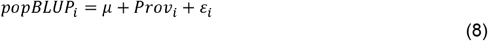

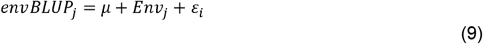

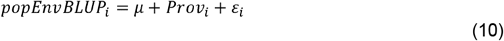

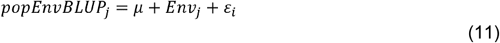

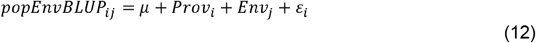

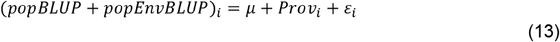

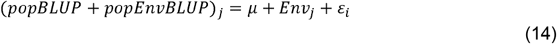

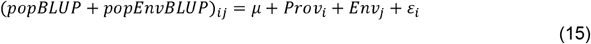

where *popBLUP*_*i*_ is the population BLUPs from model (3) as the response for model (8), *envBLUP*_*j*_ is the environment BLUPs from model (3) as the response for model (9), *popEnvBLUP*_*i*_, *popEnvBLUP*_*j*_, and *popEnvBLUP*_*ij*_ are the population × environment BLUPs from model (3) as the response for models (10-12), respectively, (*popBLUP* + *popEnvBLUP*)_*i*_, (*popBLUP* + *popEnvBLUP*)_*j*_, and (*popBLUP* + *popEnvBLUP*)_*ij*_ are the sum of the population and population × environment BLUPs from model (3) as the response for models (13-15), respectively, *µ* is the overall mean for models (8-15), *Prov*_*i*_ is the random effect of population provenance *i* in models (8, 10, 12, 13, and 15), *Env*_*j*_ is the random effect of trial environment *j* in models (9, 11, 12, 14, and 15), and *ε*_*i*_ is the random error for models (8-15). The random effects were distributed as 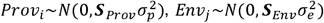, and 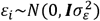. Each model was five-fold cross-validated with 20 replications. Predictive ability was estimated within each fold of each replication as the Pearson’s correlation between the observed and predicted values of the test set.

## Results

### Climatic modeling increases heritable signal for juvenile tree growth

In the “naïve” across-year phenotypic model (3), we found that population and population × environment explained a smaller proportion of the total variance in tree height (9% each) than environment, which accounted for 75% of the total variance (Table 1). Although population variance was low (9%), the highly replicated experimental design allowed for a high estimation of broad-sense heritability for tree height (H^2^ = 0.89) (Table 1). Year had an effect estimate of 13.46 ± 0.73 cm.

**Table 1.** Broad-sense heritability (H^2^) and number of levels and estimated variances for population/provenance, environment, and residual random effects from tree height models across (1980-1988) and within (1980-1983) years.

| Year | Model | Term | Estimated variance<br>± standard error | Proportion of<br>total variance | Levels | $H^2$ |
| --- | --- | --- | --- | --- | --- | --- |
| Across<br>(1980-1988) | Naive | Population | 35.67 ± 3.67 | 0.094 | 278 | 0.89 |
|  |  | Environment | 284.38 ± 39.62 | 0.747 | 107 |  |
|  |  | Pop × Env | 32.55 ± 1.42 | 0.086 | 2702 |  |
|  |  | Error | 27.91 ± 0.63 | 0.073 |  |  |
|  | Climatic | Provenance | 289.49 ± 44.30 | 0.050 | 278 | 0.96 |
|  |  | Environment | 5325.40 ± 717.78 | 0.920 | 107 |  |
|  |  | Prov × Env | 115.60 ± 4.15 | 0.020 | 2702 |  |
|  |  | Error | 59.77 ± 1.29 | 0.010 |  |  |
| 1980 | Naive | Population | 14.06 ± 1.63 | 0.333 | 278 | 0.81 |
|  |  | Environment | 19.33 ± 5.85 | 0.457 | 24 |  |
|  |  | Pop × Env | 5.78 ± 0.59 | 0.137 | 612 |  |
|  |  | Error | 3.09 ± 0.14 | 0.073 |  |  |
|  | Climatic | Provenance | 60.57 ± 9.81 | 0.362 | 278 | 0.94 |
|  |  | Environment | 96.66 ± 29.47 | 0.578 | 24 |  |
|  |  | Prov × Env | 6.84 ± 0.60 | 0.041 | 612 |  |
|  |  | Error | 3.08 ± 0.14 | 0.018 |  |  |
| 1981 | Naive | Population | 14.91 ± 1.99 | 0.169 | 278 | 0.72 |
|  |  | Environment | 56.16 ± 16.80 | 0.638 | 24 |  |
|  |  | Pop × Env | 8.50 ± 1.01 | 0.097 | 612 |  |
|  |  | Error | 8.46 ± 0.39 | 0.096 |  |  |
|  | Climatic | Provenance | 60.17 ± 11.20 | 0.160 | 278 | 0.91 |
|  |  | Environment | 297.05 ± 89.45 | 0.792 | 24 |  |
|  |  | Prov × Env | 9.57 ± 0.96 | 0.026 | 612 |  |
|  |  | Error | 8.43 ± 0.39 | 0.022 |  |  |
| 1982 | Naive | Population | 18.42 ± 3.23 | 0.073 | 278 | 0.57 |
|  |  | Environment | 191.41 ± 56.97 | 0.762 | 24 |  |
|  |  | Pop × Env | 20.31 ± 2.42 | 0.081 | 612 |  |
|  |  | Error | 20.97 ± 0.96 | 0.083 |  |  |
|  | Climatic | Provenance | 73.21 ± 16.98 | 0.069 | 278 | 0.84 |
|  |  | Environment | 938.58 ± 281.11 | 0.890 | 24 |  |
|  |  | Prov × Env | 21.55 ± 2.21 | 0.020 | 612 |  |
|  |  | Error | 20.93 ± 0.97 | 0.020 |  |  |
| 1983 | Naive | Population | 29.73 ± 6.24 | 0.042 | 278 | 0.49 |
|  |  | Environment | 582.74 ± 173.03 | 0.825 | 24 |  |
|  |  | Pop × Env | 47.06 ± 5.49 | 0.067 | 612 |  |
|  |  | Error | 46.68 ± 2.17 | 0.066 |  |  |
|  | Climatic | Provenance | 109.13 ± 30.34 | 0.041 | 278 | 0.77 |
|  |  | Environment | 2427.60 ± 724.66 | 0.922 | 24 |  |
|  |  | Prov × Env | 50.90 ± 5.07 | 0.019 | 612 |  |
|  |  | Error | 46.56 ± 2.16 | 0.018 |  |  |

We also fit a “climatic” across-year phenotypic model (4) in which the variance-covariance of the population provenance and environment random effects were modeled as the climatic similarity matrices between population provenances and trial environments, respectively. The climatic across-year model increased the proportion of the total variance in tree height explained by environment (92%) compared to the naïve across-year model (75%) (Table 1). Although population and population × environment variance were smaller (population = 5%, population × environment = 2%) than in the naïve across-year model (population = 9%; population × environment = 9%), broad-sense heritability increased (H^2^_naive_ = 0.89; H^2^_climatic_ = 0.96) (Table 1). Year had no effect on tree height in the across-year climatic model.

In the naïve within-year analysis (model 5), environmental variance increased while broad-sense heritability and population variance tended to decrease over time from 1980 (seven-year-old trees, H^2^ = 0.81, genetic variance = 33%, environmental variance = 46%) to 1983 (10-year-old trees, H^2^ = 0.49, genetic variance = 4%, environmental variance = 83%) (Table 1). In contrast, specifying the variance-covariance of the population and environment terms with climatic similarity matrices (“climatic” model 6) led to more stable estimates of broad-sense heritability (H^2^_1980 → 1983_ = 0.94 → 0.77) and population × environment variance (population × environment variance_1980→1983_ = 4 → 2%) over time (Table 1). Within-year climatic modeling (6) demonstrated an increase in environmental variance and a decrease in population variance over time (genetic variance_1980→1983_ = 36 → 4%, environmental variance_1980→1983_ = 58 → 92%) similar to that of the naïve models (Table 1).

### Climate explains environmental variance in tree height

Because the common garden experiment consisted of even-aged stands planted in the same year across trial locations, tree age (physiological growth stage) and trial environment (location-year) may have been confounded. As such, we modeled the environment BLUPs for tree height with climatic similarity between trial environments (model 9). The environmental similarity matrix, estimated from climatic and geospatial data across trial location-year environments, explained 71% of the variance in environmental BLUP estimates for tree height and had a predictive ability of r = 0.48 (Table 2). Complementarily, climatic phenotypic modeling including climatic variance-covariance matrices within (model 6) and across (model 4) years increased the variance explained by the environment term (Table 2) and removed the effect of year on tree height when compared the naïve phenotypic modeling (models 3 and 5). Combined, these results suggest that the variation in tree height estimates among trial location-year environments was largely driven by climatic variation rather than by the confounding effects of tree growth stage.

**Table 2.**
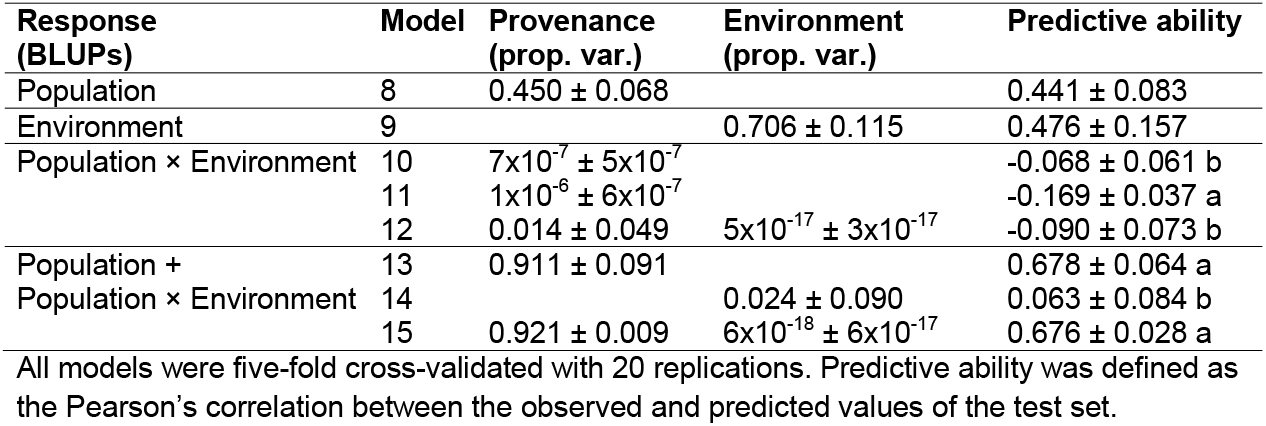
Predictive ability and proportion of total variance in population, environment, and/or population × environment best linear unbiased predictors (BLUPs) for tree height explained by population provenance and/or trial environment climatic similarity matrices.

### Provenance climate is highly predictive of inter-population and population × environment variation in tree height

The population provenance similarity matrix, estimated from historical climatic and geospatial data across population provenances, demonstrated strong population structure (Figure 1) and explained 45% of the variance in populations BLUPs for tree height (model 8), with a predictive ability of r = 0.44 (Table 2). Although population × environment BLUPs were poorly modeled (r < 0; explained variance ≤ 1%) by provenance and/or environment climatic similarity (models 10-12), provenance climatic similarity explained nearly all of the variance (>91%) in and was highly predictive (r = 0.68) of the sum of population BLUPs plus population × environment BLUPs (models 13 and 15) (Table 2). In addition, climatic phenotypic modeling (models 4 and 6) demonstrated more stable estimation of population × environment effects when compared to the naïve phenotypic models (3 and 5). These results indicate a strong signal of local adaption, where populations from provenances with similar geography/climate have similar tree growth, regardless of the trial environment in which they are observed.

**Figure 1.**
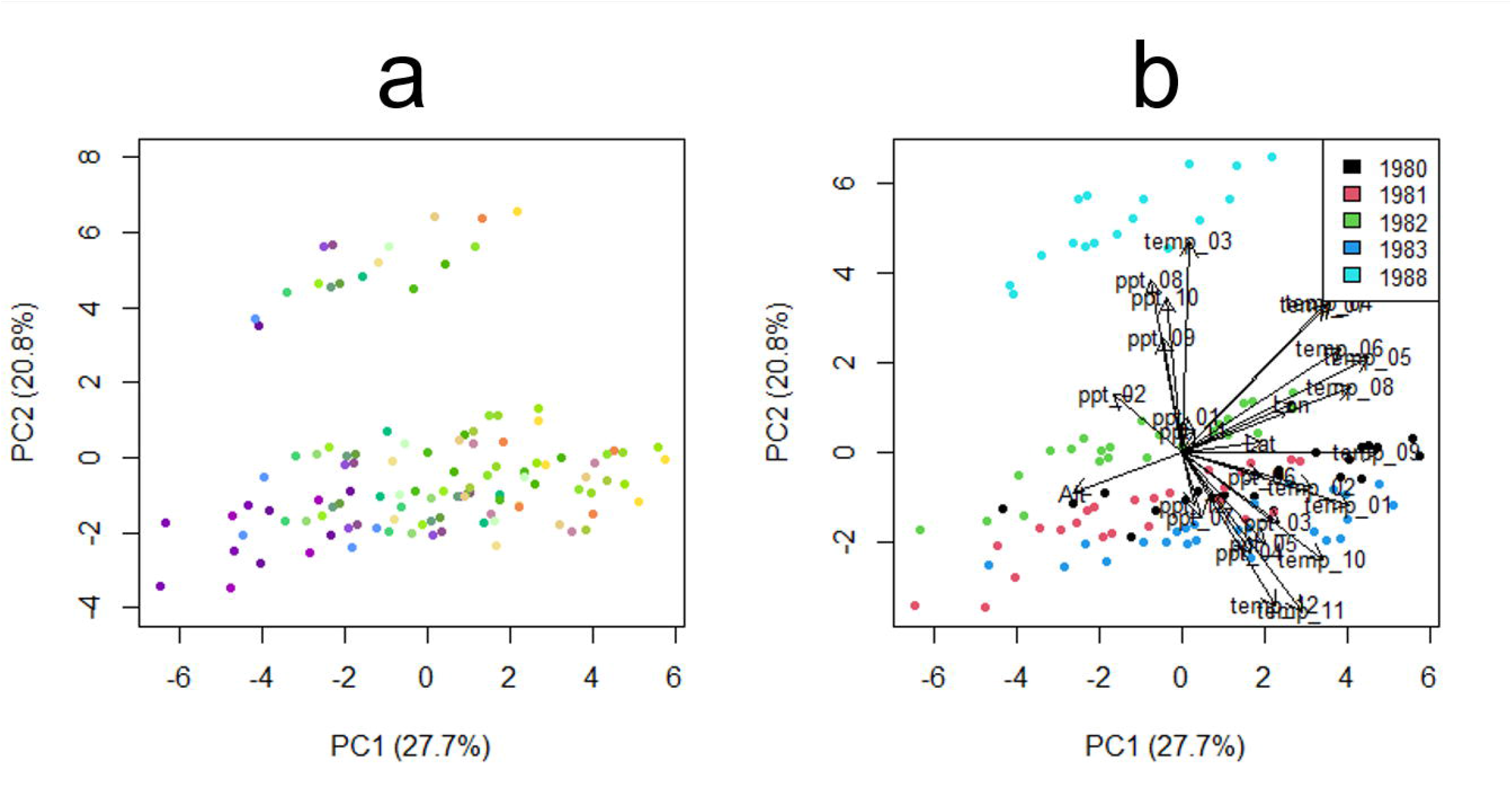
Plots of the first and second principal components (PCs) from principal component analysis (PCA) of trial environments based on geospatial and climatic data. Trial environments were labeled based on (A) a combination of altitude, longitude, and latitude and (B) year. Eigenvectors for the geospatial and weather predictors of the PCA are shown in (B).

### Climatic and geospatial variation among trial environments and population provenances

In a PCA using geospatial and weather data from trial environments (location-years), the first and second principal components (PCs) explained 28% and 21% of the variance, respectively (Figure 2). The first PC demonstrated a separation between environments based on altitude and summer/winter temperature (Figure 2). On the second PC, environments were separated by year, spring/winter temperature, and summer precipitation (Figure 2).

**Figure 2.**
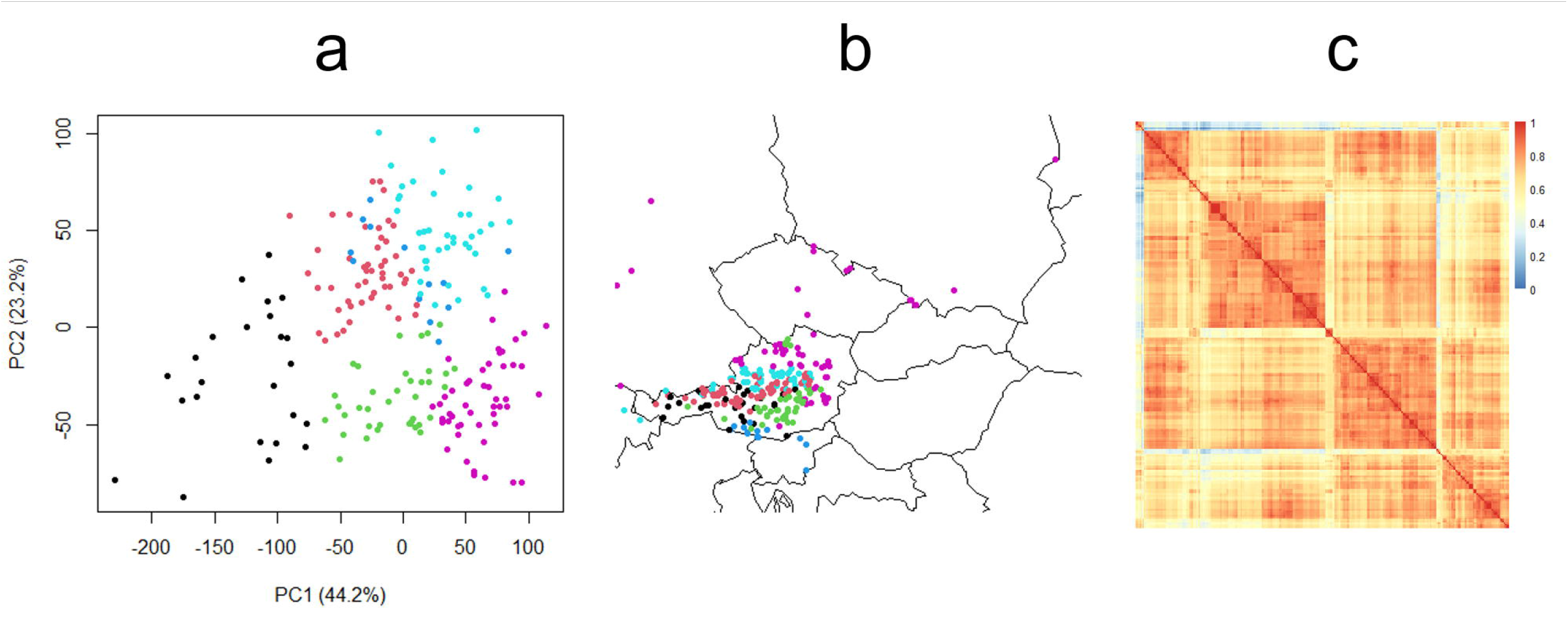
(A) Plot of the fist and second principal components (PCs) from principal component analysis (PCA) of population provenances based on geospatial and climatic data. (B) Map of population provenances. Points in (A) and (B) were labeled based on the five clusters identified by hierarchical clustering. (C) Population provenance climatic distance matrix, organized by hierarchical clustering.

We also conducted PCA for population provenances using geospatial and weather data. The first and second PCs explained 44% and 23% of the variance, respectively (Figure 1). The eigenvectors of the PCA predictor variables demonstrated that variation in PC 1 was driven by altitude, temperature, and snowfall (solid precipitation), while variation in PC 2 was driven by rainfall (liquid precipitation). Hierarchical clustering of the PCA revealed six clusters, which were geographically distributed as Carinthia, Austria and Slovenia; Styria, Austria; Lower Austria, Germany, Czech Republic, Slovakia, and Poland; and three alpine regions (Figure 1).

## Conclusions

We used agricultural statistical methods to dissect the heritable and environmental components of juvenile tree growth in Central European Norway Spruce. The highly replicated design of the common garden experiment allowed for precise estimation of genetic (population and population × environment) and environmental (trial location-year) effects. Population explained a relatively small proportion of the total variance in tree height compared to environment, similar to previous findings in this dataset (Kapeller et al. 2017; Messina et al. 2018; Washburn et al. 2020) and consistent with low estimates of population differentiation between Norway spruce populations (Androsiuk et al. 2013). However, broad-sense heritability was moderate to high and previous studies on tree height in clonal and progeny trials of Norway spruce (Hannrup et al. 2004; Chen et al. 2017) have reported similar heritability estimates. Modeling the variance-covariance of the population provenance and environment terms using climate data led to higher estimates of heritability and environmental variance and more stable estimation of heritability and population × environment for tree height across developmental time. The higher heritability estimates with climatic modeling were likely the result of increased environmental signal and subsequently decreased error variance. Tree height becomes less predictive of total biomass with age and growth habit (branching pattern etc.) may be a better indicator of tree growth after the transition between the juvenile and adult stages (Zianis et al. 2005), which may partially explain the reduction in heritability and genetic variance for tree height over time in this study. Drought conditions in Central Europe increased in severity from the beginning to the end of the trials (1983-1988), with a large historical drought event beginning in 1987 (Spinoni et al. 2015), supporting the PCA separation between the 1988 and 1981-1983 trial years reported here. In addition, the increasing environmental variance for tree height with developmental age may be partially explained by increasing drought severity across the trial period.

Although genetic marker data was not available in this dataset, we modeled inter-population relationships using historical climatic data from the population provenances. Plant and animal breeders frequently use climate and/or genetic marker data to model relationships among individuals and/or environments in order to improve the prediction accuracy of phenotypes in new (genetically related) material and even in new environments (Heslot et al. 2014; Jarquín et al. 2014; Crossa et al. 2016; Chen et al. 2017; Messina et al. 2018; Bustos-Korts et al. 2019; Gillberg et al. 2019; Washburn et al. 2020; Costa-Neto et al. 2021). Previous studies have demonstrated that local adaptation is prevalent in tree species, including Norway spruce, and that ancestral environment is a strong predictor of growth phenotypes in Norway spruce (Alberto et al. 2013; Berg and Coop 2014; Aitken and Bemmels 2016; Milesi et al. 2019). We found that climatic relationships among population provenances (a) were highly predictive of population + population × environment tree height variation and (b) demonstrated population structure nested within geography, indicating that these populations have strong local adaptation (Aitken and Bemmels 2016). Our findings complement previous results from ecological modeling of the dataset, where within-population variation was driven by population provenance temperature (Kapeller et al. 2017). The modeling used in this study results in refined tree growth estimates for population and population × environment, which could be used to identify potential sources of material adapted to new environments.

## Author contributions

Conceptualization, LM; Data curation, LM; Formal analysis, LM; Funding acquisition, KS; Methodology, LM; Project administration, LM; Software, LM; Validation, LM; Visualization, LM; Writing-original draft, LM; Writing-review and editing, LM, KS

